# Acute Paternal Immune Activation Shapes Embryonic Development and Protects Offspring from Viral Infection

**DOI:** 10.64898/2026.03.06.710215

**Authors:** Taylor Miller-Ensminger, Ariana D. Campbell, Elizabeth A. Kennedy, Madeline N. Lamonica, Cecelia N. Murphy, Grace S. Lee, Mia Weller, Montserrat C. Arreguin, Madeline S. Merlino, Seble G. Negatu, Peter Hewins, Colin C. Conine, Kellie A. Jurado

**Author notes:** These authors contributed equally (Ariana D. Campbell, Elizabeth A. Kennedy, Madeline N. Lamonica).

## Abstract

The evolutionary arms race between host and pathogen is traditionally framed as a multigenerational paradigm, where beneficial host immune adaptations accrue in a population over time through DNA sequence variation. If pathogens exposure alters epigenetic information inherited by offspring through paternal gametes, advantageous immune traits could emerge within a single generation. We show that acute paternal immune activation (PIA) induces immune signaling in the male reproductive tract and remodels the sperm small RNA profile. Using IVF, we show that altered sperm small RNAs from immune-activated males reprograms gene expression in preimplantation embryos, eliciting sex-specific transcriptional responses. This transcriptional program is reflected in a striking inherited phenotype, as male offspring sired by PIA males exhibit enhanced survival following lethal viral challenge. These findings identify sperm small RNAs as a molecular mechanism that links PIA to embryonic development and offspring immunity, providing a comprehensive framework for how PIA shapes inherited pathogen protection.

## INTRODUCTION

Over the last decade, our understanding of paternal influences on offspring has expanded dramatically. Once thought to only contain genetic information encoded in DNA, it is now recognized that sperm carry diverse forms of epigenetic information that shape embryonic development and offspring physiology.^1–3^ For example, environmental stressors experienced by parents remodel epigenetic information in germ cells that is passed to subsequent generations, causing observable offspring phenotypes.^1,2^ Through this mechanism, termed intergenerational non-genetic inheritance, ancestral environments can alter epigenetic information that is transmitted to progeny. This enables adaptations to arise in a single generation, independently of alterations to the DNA sequence.

Mediators of epigenetic inheritance include DNA methylation, histone post-translational modifications, and small regulatory RNAs.^1,2,4–7^ While each of these molecules are present in sperm and have the potential to carry heritable information, small RNAs have been causally demonstrated to transmit phenotypes from male parents to offspring in mammals.^2,8^ The paternal small RNA profile is initially established during spermatogenesis in the testes and then subsequently altered during epididymal transit by interacting with epididymosomes, extracellular vesicles released by the epididymal epithelium.^3,9,10,11^ These vesicles deliver cargo, including small RNAs, to sperm in the epididymal lumen. Importantly, two classes of small RNAs acquired during epididymal transit, microRNAs (miRNAs) and tRNA-Derived RNAs (tDRs), have been shown to directly regulate embryonic gene expression and preimplantation development.^3,12,13^ Although sperm small RNAs are highly influenced by the paternal environment, how these molecules transfer lasting non-genetically inherited phenotypes in offspring remains undetermined.^2^

Though a variety of experimental paradigms have established paternal intergenerational non-genetic inheritance in mammals, most studies have focused on evaluating the impacts of diet, toxin, and stress exposure on epigenetic information in sperm and offspring outcomes.^1,2,14,15^ Interestingly, many studies report sex-specific phenotypes,^2,16^ suggesting that impact of epigenetic inheritance is tied to offspring sex. Collectively, these studies have provided substantial insight into how paternal stressors can alter offspring metabolism and behavior. However, few mammalian studies have explored the influence of paternal intergenerational non-genetic inheritance on offspring immunity. In contrast, a phenomenon referred to as transgenerational immune priming (TGIP) is well established in non-mammalian models. In models of TGIP, parental pathogen exposure enhances offspring resistance to homologous or heterologous infections.^17–19^

Recent research has begun to unravel the interplay between immune activation in parents and offspring phenotypes in mammalian models. Specifically, maternal immune activation (MIA) during gestation, via both lipopolysaccharide (LPS) and Polyinosinic:polycytidylic acid (Poly(I:C)), can have lifelong impacts on offspring immunity, behavior, and cognitive outcomes.^20–23^ Importantly, during MIA, developing fetuses are directly exposed to the maternal inflammatory environment. In contrast, the impacts of paternal immune activation (PIA), an event that occurs prior to conception, on offspring are only beginning to emerge. The few studies exploring PIA have implemented an array of microbial or immune stimulatory triggers and have primarily focused on assessing offspring behavior outcomes.^24–26^ While there is an established link between PIA and offspring behavior, the relationship between PIA and offspring immunity is unclear. Limited evidence suggests that PIA alters offspring immune responses, primarily in systemic cytokine response,^25–27^ though it is unclear if PIA protects offspring in an infection context.^28,29^ Additionally, prior studies focused on the effects of PIA at 4+ weeks post-stimulation, overlooking the impacts of immune stimulation immediately prior to conception. Together, these studies reveal a fundamental gap in our understanding of how acute paternal immune activation prior to conception programs offspring immune function and infection outcomes.

Here, we evaluate the short-term consequences of PIA on sperm small RNAs and offspring immunity. We explore the short-term consequences of PIA on sperm small RNAs using two models of acute viral immune stimulation, Zika virus infection and Poly(I:C) treatment. We then define the broader impacts of acute immune stimulation on the male reproductive tract, revealing a connection between cytokine signaling and establishment of the sperm small RNA profile in the epididymis. Using single-embryo RNA sequencing, we show that PIA-remodeled sperm RNAs reprogram early embryonic gene expression, revealing sex-specific transcriptional programs. Finally, we demonstrate that male offspring sired by PIA males exhibit strikingly enhanced survival and dramatically improved infection outcomes when challenged with a lethal dose of influenza A. Our study demonstrates a distinct link between acute paternal pathogen exposure and altered offspring immunity, providing evidence that epigenetic modifications contribute to host responses during host-pathogen dynamics.

## RESULTS

### Acute viral stimulation drives remodeling of the sperm small RNAs via the cauda epididymis

We first aimed to determine if acute pathogen exposure would alter the sperm small RNA profile during epididymal transit. In mice, sperm traverse from the testis through the caput, corpus, and cauda epididymis, completing epididymal transit over the course of ∼10 days.^30,31^ To capture the effects of PIA during this critical period, we applied two models of acute PIA in which sperm are only exposed to immune activation during epididymal transit (Figure 1A). We used an immunocompetent mouse model of Zika virus infection (ZIKV) to first examine the consequences of viral infection in the male reproductive tract.^32^ In this model, we observed high levels of viral RNA and infectious virus in the epididymis of ZIKV-infected males at 5 days post-infection, with clearance by 20 days post-infection (Supplemental Figure 1A and B). To assess how acute epididymal infection altered the small RNA profile of sperm that were transiting during this period, we harvested mature sperm from the cauda epididymis at 5 days post-infection for small RNA-seq. Interestingly, we found that mice infected with ZIKV had a remodeled sperm small RNA profile, with alterations primarily to microRNAs (miRNAs) and tRNA-Derived RNAs (tDRs) (Figure 1B, Supplemental Table 1).

**FIGURE 1:**
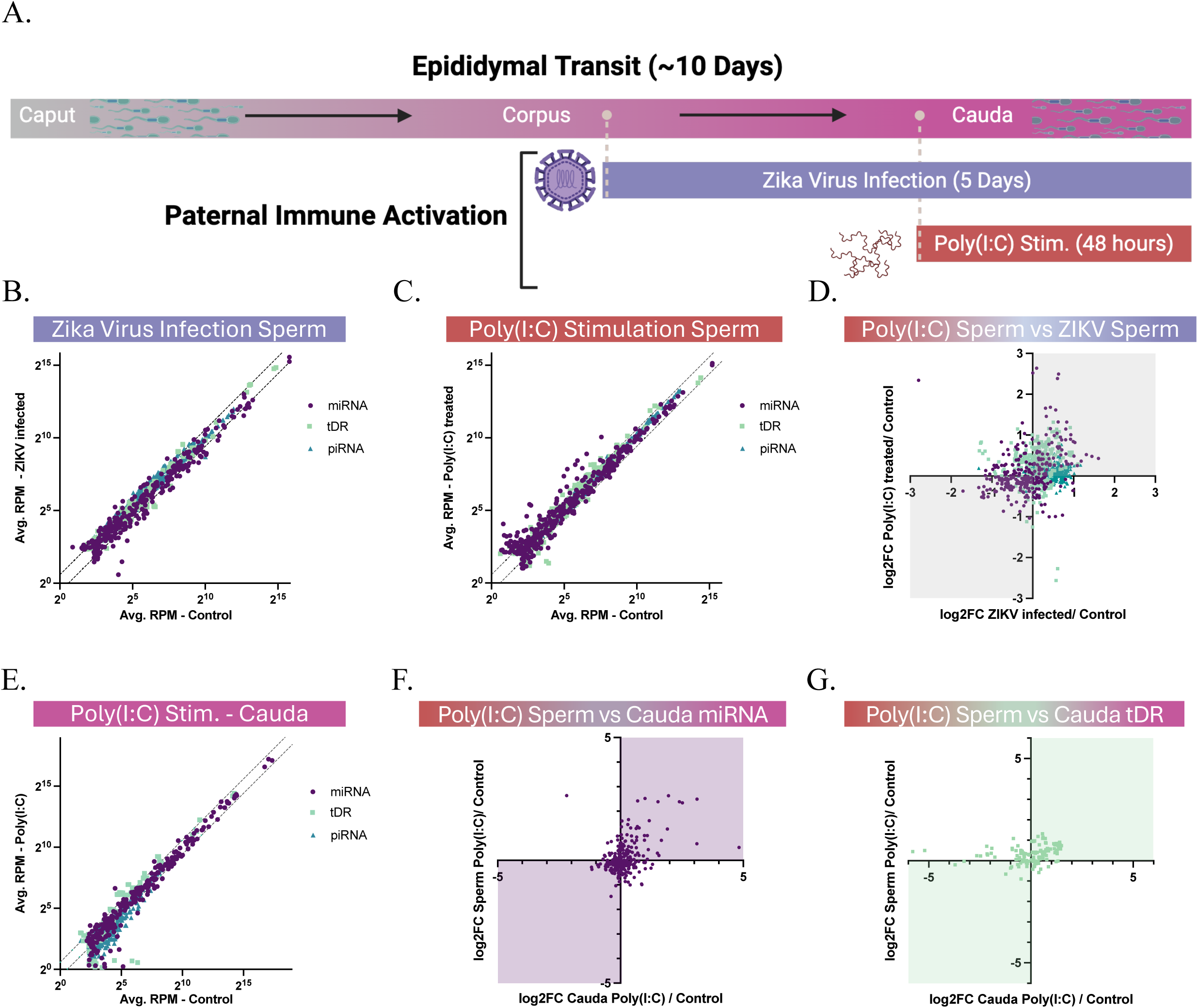
Acute viral stimulation drives remodeling of the sperm small RNAs via the cauda epididymis. A. Overview of epididymal maturation and timeline for models of paternal immune activation. B. 13-week-old male mice were infected with 10^5 PFU ZIKV DAKAR MA. Sperm were harvested 5 days later for small RNA-seq. MicroRNAs = miRNA (purple), tRNA-Derived RNAs= tDR (green), Piwi-interacting RNAs = piRNA (blue). Lines indicate 1.5-fold change. rpm = reads per million. C. 13-week-old male mice were treated with Poly(I:C). Sperm were harvested at 48 hours post treatment for small RNA-seq. D. Correlation plot showing the shared alterations to the sperm small RNA profile during Poly(I:C) treatment and ZIKV infection. Concordant changes in expression are highlighted in grey boxes. Pearson Correlation Coefficient = .1135, p value =0.0002. E. Caudal tissues were harvested at 48 hours post-Poly(I:C)-treatment for small RNA-seq. F. Correlation plot showing overlaps in changes to the miRNA profile in sperm and in the cauda after Poly(I:C) treatment. Concordant changes in expression are highlighted in purple boxes. Pearson Correlation Coefficient = 0.3791, p value < 0.0001 G. Correlation plot showing overlaps in changes to the tDR profile in sperm and the cauda after Poly(I:C) treatment. Concordant changes are highlighted in green boxes. Pearson Correlation Coefficient = 0.2408, p value = 0.0025.

We next asked whether the immune response to infection drove the observed changes to the sperm small RNA profile. To do this, we implemented a second PIA model in which male mice were treated with Poly(I:C), a synthetic viral mimetic. As Poly(I:C) rapidly drives proinflammatory responses (Supplemental Figure 1C), we harvested mature sperm from the cauda epididymis of male mice at 48 hours post-treatment to measure the acute response to stimulation (Figure 1A). Similar to the changes induced by ZIKV infection, the small RNA profile established with Poly(I:C) stimulation altered many miRNAs and tDRs (Figure 1C, Supplemental Table 1). We found strong, concordant changes in miRNAs across both models (r = 0.2816, p-value <0.0001), with a less dramatic correlation of tDRs between immune stimulatory conditions (r = 0.08714, p-value = 0.1130) (Figure 1D). These parallel alterations between the two models of acute PIA suggest that remodeling of the sperm small RNA profile is primarily driven by immune response to viral stimulation, rather than viral replication.

Epididymosomes, extracellular vesicles secreted by the epididymis, shuttle cargo from the epididymal epithelial cells to sperm, remodeling the sperm small RNA profile.^10,33^ To probe whether changes in epididymal tissue small RNA profiles were correlated, and potentially responsible for the alterations in sperm small RNAs, we sequenced small RNAs from the cauda epididymis of Poly(I:C) treated mice. This analysis revealed that the cauda epididymis is responsive to Poly(I:C) treatment, resulting in differential expression of many miRNAs, tDRs, and Piwi-interacting RNAs (piRNAs) (Figure 1E). Intriguingly, we found that alterations to the miRNA (Figure 1F, r = 0.3791, p-value < 0.0001) and tDR (Figure 1G, r = 0.2408, p-value = 0.0025) profiles in the caudal tissue closely matched the changes detected within the sperm. This suggests that Poly(I:C) treatment alters small RNA expression in the cauda, driving the changes observed in the sperm of treated mice. Taken together, our data reveal that acute viral stimulation, with or without active virus infection, can reshape the sperm small RNA profile and uncovers a role for the cauda epididymis in communicating epigenetic information to sperm.

### The cauda epididymis exhibits a proinflammatory response to acute PIA

We next assessed whether PIA directly impacts gene expression throughout the male reproductive tract. Sperm are generated in the testes during spermatogenesis. Sperm then undergo maturation during epididymal transit which begins in the caput epididymis, continues through the corpus epididymis, and finally reaches the cause epididymis where mature sperm is stored prior to mating (Figure 2A). We harvested the testes and three regions of the epididymis—the caput, corpus, and cauda—at 48 hours post-Poly(I:C) treatment for mRNA sequencing. We found altered gene expression in all tissues, with the caput and cauda epididymis being the most responsive (Figure 2B, Supplemental Table 2). When overlaps in gene expression changes were assessed, only 17 upregulated genes were shared across the four tissues assessed, with two shared downregulated genes (Figure 2C, Supplemental Table 3). Neighboring regions of the epididymis shared the most synonymous changes in gene expression, with the corpus and the cauda sharing the most alterations (Figure 2C). We next assessed which biological processes were associated with the changes in gene expression. Throughout the epididymis, we found enrichment in genes driving GO terms relating to immune and pathogen response pathways, with the most pronounced changes in the cauda epididymis (Supplemental Figure 2, Supplemental Table 4). This indicates the cauda epididymis is particularly responsive to Poly(I:C) stimulation. Many of the genes driving these GO terms were chemokines and interferon stimulated genes, thereby revealing a proinflammatory response to treatment. When we compared gene expression, with and without Poly(I:C) stimulation, across the testes and epididymal regions, we found that the proinflammatory response is largely limited to the epididymis, with the cauda showing the highest proinflammatory signature (Figure 2D).

**FIGURE 2:**
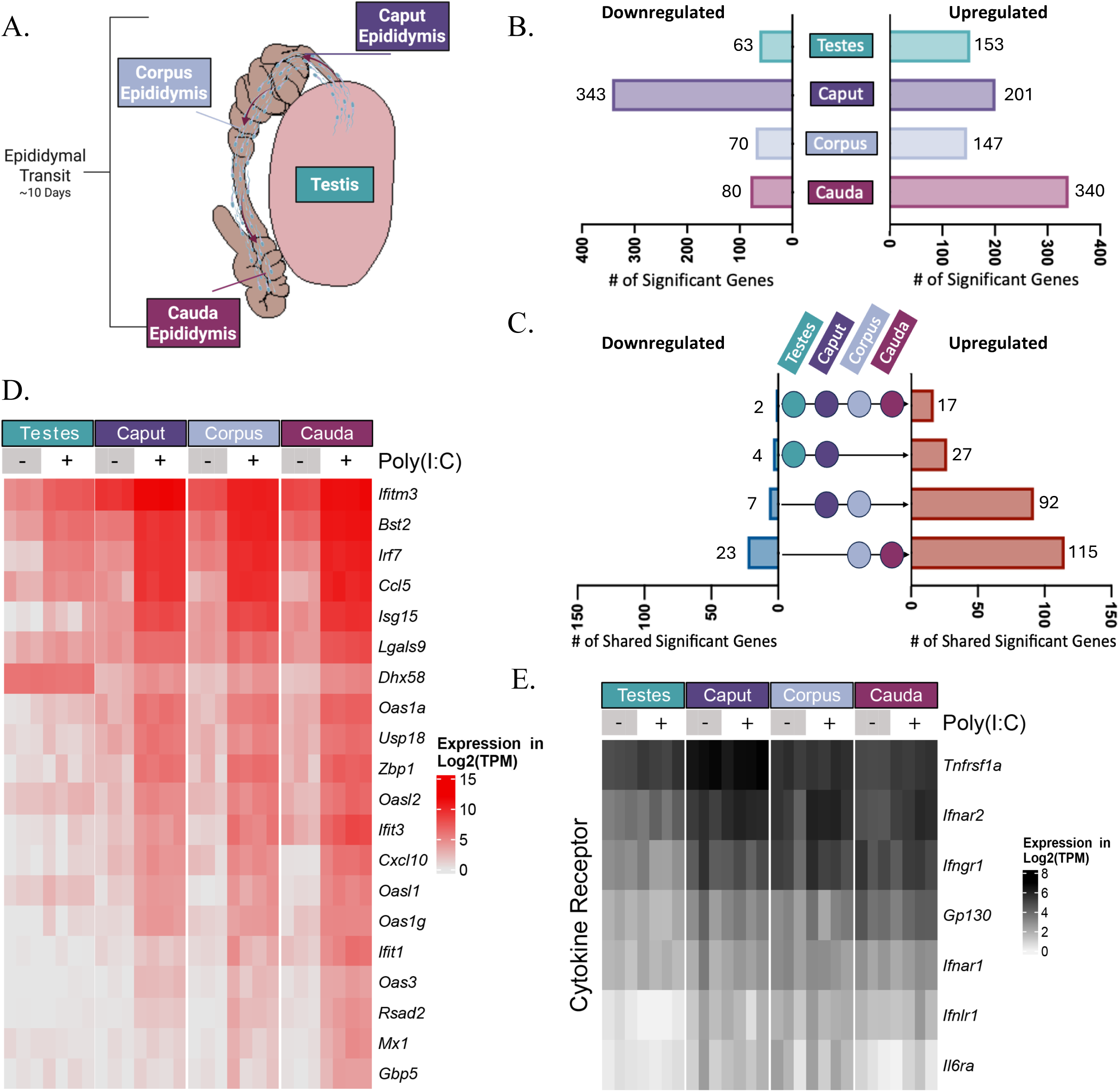
The cauda epididymis exhibits a proinflammatory response to PIA. 13-week-old male mice were treated with Poly(I:C). Testis and epididymal tissues were harvested at 48 hours post treatment for bulk mRNA-seq. A. Overview of the male reproductive tissues. B. Number of significantly up- and downregulated genes during Poly(I:C)-treatment by tissue. n = 4 Poly(I:C) treated mice, n = 3 control (PBS treated) mice. Differential expression determined by DESeq2 and significance determined by p adj < 0.05 and log_2_FC > 0.58 or log_2_FC < -0.58 (1.5 fold-increase or decrease). C. Overlapping changes in gene expression between neighboring tissues of the male reproductive tract during Poly(I:C) treatment. D. Expression of proinflammatory and immune genes driving upregulated GO term analysis in the cauda epididymis (defined in S2) within all tissues of the male reproductive tract in Poly(I:C) treated and control mice. E. Expression of cytokine receptors in the male reproductive tract in Poly(I:C) treated and control mice.

We next sought to understand why the epididymis is particularly responsive to PIA. Because Poly(I:C) treatment triggers systemic proinflammatory cytokine signaling, we evaluated the expression of cytokine receptors in the testes and epididymal regions. Here, we found that regardless of treatment state, the epididymis highly expresses tumor necrosis factor receptor superfamily 1a (*Tnfrsf1a*), interferon alpha/beta receptor subunit 2 (*Ifnar2*), and interferon gamma receptor 1 (*Ifngr1*) (Figure 2E). High expression of these genes as well as moderate expression of other receptors (*Gp130*, *Ifnar1*), suggests that the epididymis is primed to respond to systemic cytokine signaling. Altogether, these data suggests that the male reproductive tract, particularly the cauda epididymis, is primed and highly responsive to systemic responses to infection.

### PIA alters early embryonic gene expression

Given the impact of PIA on the sperm small RNA profile, we sought to determine if these changes influence embryogenesis. Sperm from PIA (Poly(I:C)-treated) or control males was collected at 48 hours post-treatment and used for in vitro fertilization (IVF) with eggs from naïve females (Figure 3A, Supplemental Figure 3A). Embryos were collected at the 4-cell and morula developmental stages for single embryo mRNA-seq. At the 4-cell and morula stages, embryos fertilized with sperm from PIA males displayed distinct changes in gene expression from control-fertilized embryos (4-cell: 84 upregulated and 74 downregulated DEGS; morula: 43 upregulated and 73 downregulated DEGs) (Figure 3B and 3D, Supplemental Table 5).

**FIGURE 3.**
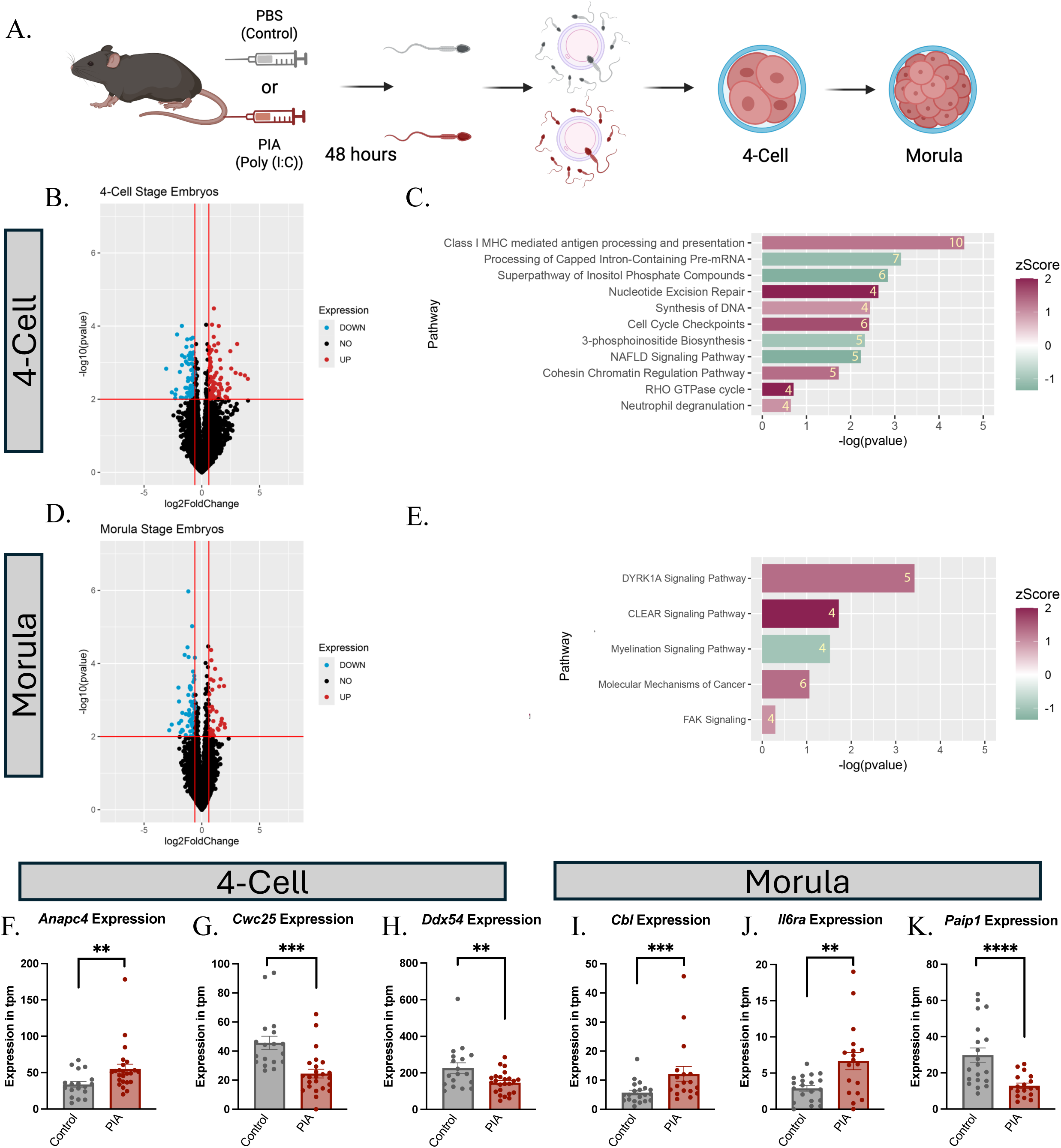
PIA alters early embryonic gene expression. A. Naïve eggs were fertilized with sperm from PIA (Poly (I:C) treated) or Control (PBS treated) males at 48 hours post-treatment. Developing embryos were harvested at the 4-cell and morula developmental stages for single embryo RNA-seq. B. Volcano plot showing differential gene expression in 4-cell stage embryos. Red lines indicate significance thresholds (−0.58 > log_2_FC or log_2_FC > 0.58 and p value < 0.01). Blue dots = downregulated, red dots = upregulated. n = 24 PIA embryos, 18 control embryos. C. Ingenuity Pathway Analysis (IPA) of differentially expressed genes in 4-cell stage embryos. Inset yellow numbers indicated number of genes involved pathway term. D. Volcano plot showing differential gene expression in morula stage embryos. E. IPA of differentially expressed genes in morula stage embryos. n = 18 PIA embryos, 20 control embryos. F-H. Examples of differentially expressed genes in 4-cell stage embryos. I-K. Examples of differentially expressed genes in morula stage embryos. F-K. P values calculated by DESeq2, P values: ** = p value < 0.01, *** = p value < 0.001, **** = p value < 0.0001.

To probe the causal pathway relationships corresponding with these changes in gene expression, we performed Ingenuity Pathway Analysis (IPA) to explore up- (pink) and downregulated (green) pathways (Figure 3C and 3E, Supplemental Table 6). While most of the differentially regulated pathways related to regulatory processes, we interestingly found that two immune pathways, Class I MHC mediated antigen processing and presentation and neutrophil degranulation, were upregulated in 4-cell embryos (Figure 3C). There were fewer altered pathways in morula embryos, and those that were affected related primarily to cellular proliferation and neuronal development and function (Figure 3E).^34–37^ In the 4-cell embryos, we examined expression of genes driving pathways in our IPA analysis. In PIA IVF embryos, we saw an increase in expression of *Anapc4,* a gene involved in cohesin chromatin regulation, cell cycle checkpoints, synthesis of DNA, and Class I MHC processing and presentation (Figure 3F). Additionally, *Cwc25,* a gene involved in processing of capped intron-containing pre-mRNA and *Ddx54,* an RNA helicase, were downregulated in PIA fertilized 4-cell. (Figure 3G and 3H). In the morula stage embryos, we observed an overall increase in expression of a cancer-related gene, *Cbl,* and a gene encoding a proinflammatory cytokine receptor, *Il6ra,* in PIA fertilized embryos compared to controls (Figure 3I and 3J). In addition to altered expression of the IL-6 receptor, we found that *Paip1*, an immune regulatory gene,^38^ had significantly lower expression in PIA fertilized embryos (Figure 3K). Taken together, these data show that fertilization by PIA sperm modulates early embryonic gene expression, including expression of immune related genes.

### PIA alters embryonic gene expression in a sex-dependent manner

As many paradigms of intergenerational non-genetic inheritance in mammals show sex-based outcomes in offspring,^2,16^ we explored whether early embryonic gene expression followed sex-specific patterns. Using the single embryo RNA-seq data we generated, embryos were separated as females and males based on expression of *Xist* (only expressed in females) and *Eif2s3y* (Y-chromosome gene only expressed in males), respectively.^39^ In 4-cell embryos, gene expression was altered for a greater number of genes in female embryos (Figure 4A and 4C, Supplemental Table 7) compared to male embryos (Figure 4B and 4C, Supplemental Table 7). In contrast, in morula stage embryos, more genes were significantly altered in male embryos than female embryos (Figure 4D-F). This suggests alterations in gene expression due to PIA fertilization are linked to sex and developmental stage. Interestingly, the majority of DEGs at both the 4-cell and morula developmental stages were found on autosomes, demonstrating that differences in mRNA expression between male and female embryos is not due to the intrinsic features of embryo sex (Supplemental Figure 3B-D, Supplemental Table 8). To further explore sex-specific PIA induced alterations in gene expression, we next assessed expression levels in our previous genes of interest based on sire and embryo sex. Here, we found three different trends in our data. First, *Anapc4* and *Il6ra* are significantly more expressed in male embryos fertilized by PIA males, but not significantly altered in female embryos (Figure 4G and 4K). *Cwc25* and *Ddx54* follow the converse pattern with reduced expression in female embryos fertilized by PIA males (Figure 4H and 4I). Finally, fertilization by PIA males alters expression of *Cbl* and *Paip1* in both male and female embryos (Figure 4J and 4L). Taken together, these data show that PIA induces sex-specific changes within the early embryo that are not limited to genes on the sex chromosomes.

**FIGURE 4.**
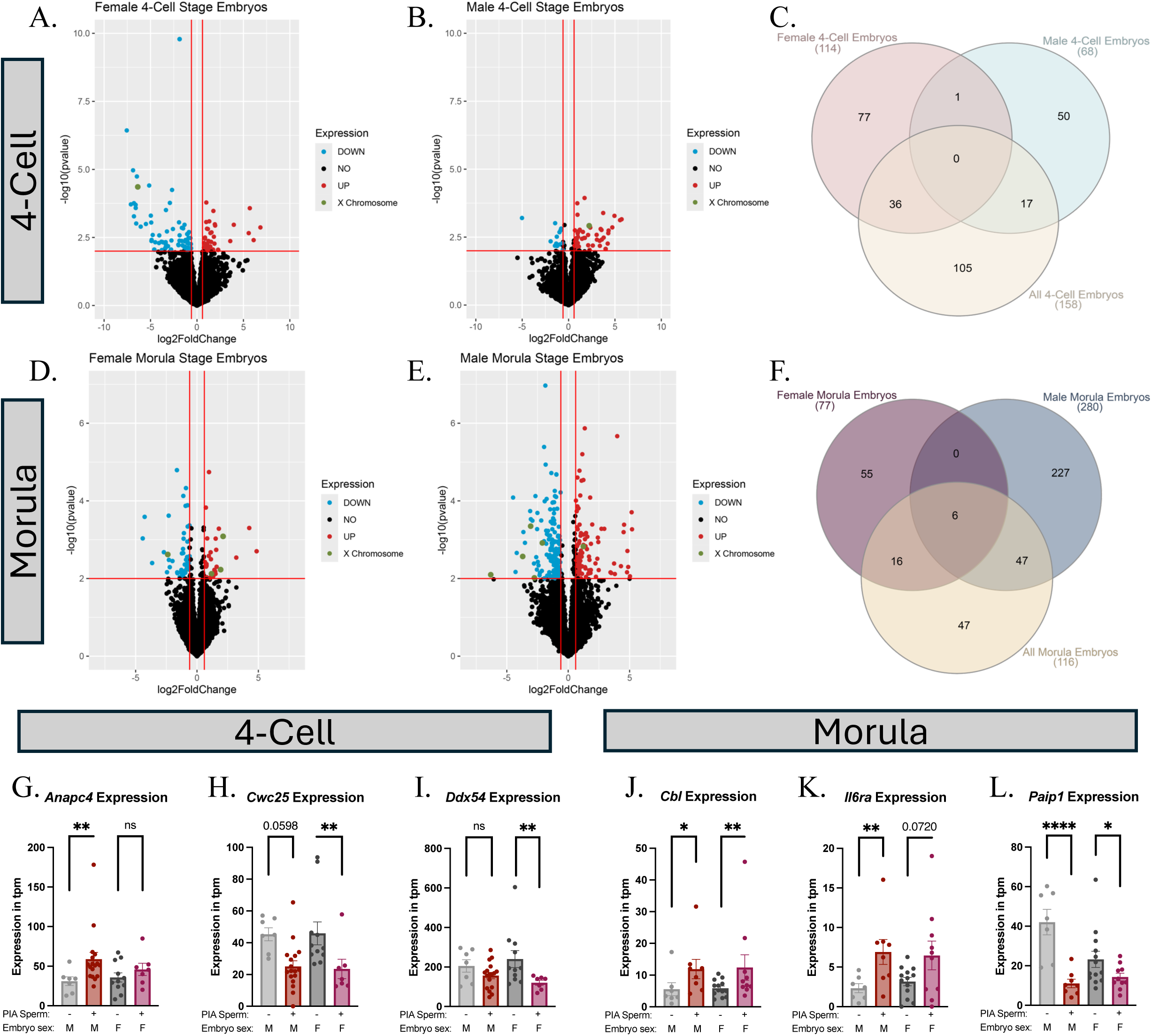
PIA alters embryonic gene expression in a sex-dependent manner. Naïve eggs were fertilized with sperm from PIA (Poly(I:C) treated) or Control (PBS treated) males at 48 hours post-treatment. Developing embryos were harvested at the 4-cell and morula developmental stages for single embryo RNA-seq. Embryo sex was determined by the expression of *Xist* (female) and *Eif2s3y* (male). A. Volcano plots showing differential gene expression in female 4-cell stage embryos. n = 7 PIA embryos, 11 control embryos. B. Volcano plot showing differential gene expression in male 4-cell stage embryos. n = 17 PIA embryos, 7 control embryos. C. Overlapping changes in gene expression between male, female, and all 4-cell stage embryos. D. Volcano plots showing differential gene expression in female morula stage embryos. n = 10 PIA embryos, 13 control embryos. E. Volcano plots showing differential gene expression in male morula stage embryos. n = 8 PIA embryos, 7 control embryos. F. Overlapping changes in gene expression between male, female, and all morula stage embryos. A,B,D,E. Red lines indicate significance thresholds (−0.58 > log_2_FC or log_2_FC > 0.58 and p value < 0.01). Blue dots = downregulated, red dots = upregulated, green dots = genes encoded on the X chromosome. G-I. Examples of differentially expressed genes in 4 Cell stage embryos. J-L. Examples of differentially expressed genes in Morula stage embryos. G-L. P values calculated by DESeq2, P values: ns = p value > 0.05 * = p value < 0.05, ** = p value < 0.01, **** = p value < 0.0001.

### PIA sperm RNAs are sufficient to alter embryonic gene expression

Next, we sought to determine whether RNAs from the sperm of PIA treated males were responsible for altering gene expression in the early embryo. To test this, total and small RNA were isolated from the sperm of PIA (Poly(I:C)) treated males at 48 hours post-treatment and microinjected into embryos fertilized by control sperm (Figure 5A, Supplemental Figure 4A). Nuclease free water (NFW)-injected embryos and PIA IVF embryos served as negative and positive controls, respectively, and morula-stage embryos were collected for single embryo mRNA-seq. Strikingly, when comparing the embryonic gene expression established by PIA fertilization and total RNA injections, we found a significant correlation in altered gene expression patterns (r = 0.5398, p-value <0.0001) (Figure 5B). Moreover, many of the DEGs identified in our original gene expression analysis of embryos fertilized by PIA sperm (Figure 3D), followed the same pattern of differential expression in total RNA injected embryos (Figure 5B, Supplemental Figure 4B). Similarly, injection of the sperm small RNAs from PIA males also led to changes in gene expression that mimicked those seen when embryos were fertilized with sperm from PIA males (r = .4391, p-value <0.0001) (Figure 5C) and induced similar expression patterns in original gene expression analysis of embryos fertilized by PIA sperm (Figure 3D, Figure 5C, Supplemental Figure 4C). Collectively, these data indicate that total and small RNAs from the sperm of PIA males are sufficient to alter embryonic gene expression in PIA fertilized embryos.

**FIGURE 5.**
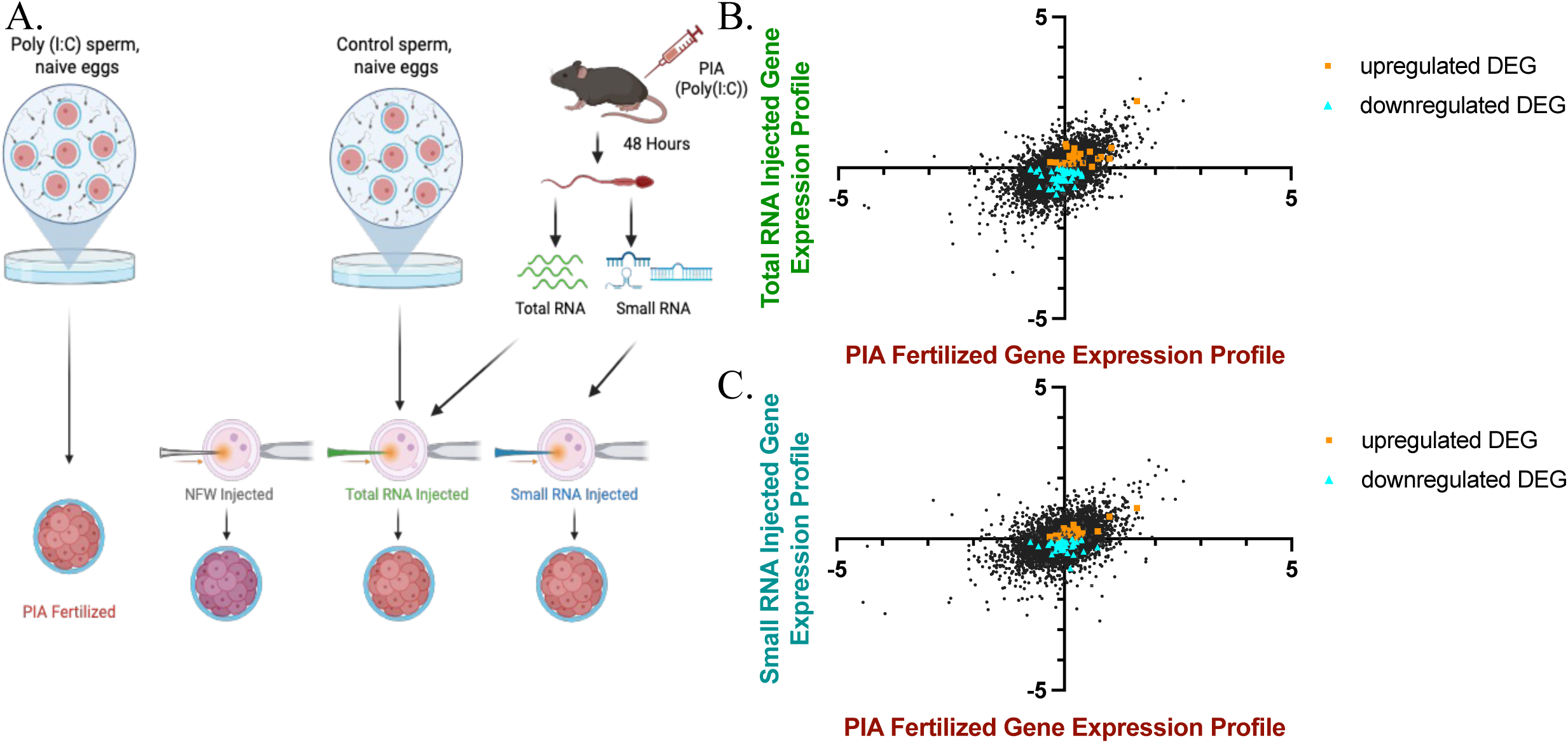
PIA sperm RNAs are sufficient to alter embryonic gene expression. A. Overview of experimental design used for the creation of morula stage embryos for single embryo sequencing. B. Comparison of profile established by total RNA injection to PIA fertilized profile, both profiles normalized to NFW embryos. Pearson Correlation. r = .5398, p value <0.0001. C. Comparison of profile established by small RNA injection to PIA fertilized profile, both profiles normalized to NFW embryos. Pearson Correlation. r = .4391, p value <0.0001. B and C. Orange squares indicate genes that were upregulated in original embryo sequencing experiments (Figure 3) and total or small RNA injection embryos. Blue triangles indicate genes that were downregulated in original embryo sequencing experiments (Figure 3D) and total or small RNA injection embryos. Only concordant changes from Figure 3 are shown.

### Paternal immune activation protects male offspring from lethal viral infection

Paternal stressors are often linked to offspring outcomes, for example, paternal diet has been found to alter offspring metabolism.^2^ Therefore, we next aimed to determine if acute PIA altered offspring immunity. We generated offspring using our Poly(I:C) model of acute PIA. Male mice were treated with Poly(I:C) 48 hours prior to mating to induce PIA which was confirmed by increased levels of serum IL-6, (Figure 6A-B). Naïve females were then mated to PIA males at 48 hours post-treatment and offspring from these matings were infected with an adult lethal dose of influenza A strain PR8 at 23 days old. Following infection, offspring were monitored daily for changes in weight, temperature, and clinical signs for 14 days. Fascinatingly, we found that male offspring, but not female offspring, sired by PIA males had a strikingly higher survival rate of 71% compared to offspring sired by control males which ranged from 15-18% (Figure 6C and 6G, Supplemental Figure 5A and 5B). Further, PIA sired male offspring better maintained core body temperature, lost less weight, and had fewer clinical signs associated with infection, as compared with control sired males (Figure 6D-F, Supplemental Figure 5C, 5E, and 5G). Conversely, female offspring showed no difference in infection severity, regardless of sire (Figure 6H-J, Supplemental Figure 5D, 5F, and 5H). Next, we explored whether protection in male offspring sired by PIA males was correlated with lower viral burden in the lung. Lungs were harvested from infected mice at 4- or 6/7-days post infection and infectious viral burden was measured through TCID50 assay. Interestingly, all mice showed similar levels of virus in the lung at both collection timepoints (Supplemental Figure 5I and 5J). These data suggest that differences in survival and infection severity are not linked to better control of viral replication or clearance, but rather suggests differential immune response activation in the offspring. Taken together, our results show that PIA influences offspring susceptibility to infection, conferring protection to male offspring during a lethal influenza A challenge.

**FIGURE 6.**
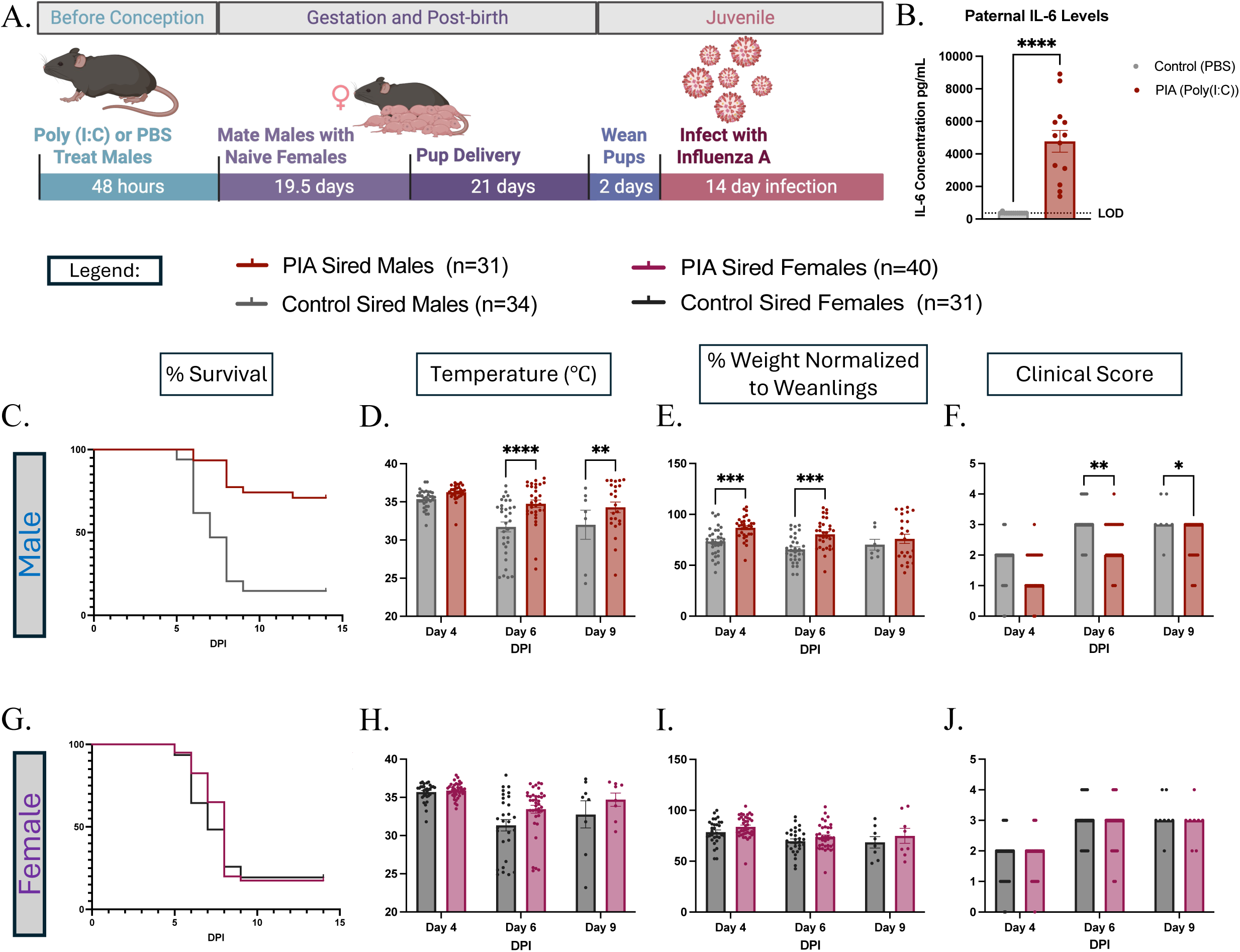
PIA confers resistance to viral infection in male offspring. A. Overview of treatment of PIA males and mating timeline for offspring creation and infection. Offspring sired by PIA (Poly(I:C) treated) and Control males were infected intranasally with 250 TCID50 IVA PR8 at 23 days old. B. Male parents were treated with Poly(I:C) or PBS. At 6 hours post treatment, serum was collected and circulating IL-6 levels were measured by ELISA. Mann-Whitney test. p value < 0.0001. LOD indicates limit of detection (365 pg/mL) C. Survival of male offspring by sire condition. Mantel-Cox test, p value < 0.0001. D-F. Clinical outcomes associated with influenza infection in male offspring. Days 4,6, and 9 are shown. Weights are normalized to uninfected litters. Sidak’s multiple comparisons test. * = p value < 0.05, ** = p value < 0.01, *** = p value < 0.001. G. Survival of female offspring by sire condition. Mantel-Cox test, p value = 0.4984. H-J. Clinical outcomes associated with influenza infection in female offspring. B-J. Graphs are representative of 15 PIA sired litters and 14 Control sired litters from 14 and 13 males respectively.

## DISCUSSION

In this study, we define the impacts of acute PIA on the male reproductive tract, sperm small RNAs, and PIA sired offspring development. Using two models of acute viral immune stimulation, we demonstrate that PIA rapidly remodels the sperm small RNA profile as quickly as 48 hours post-immune stimulation. Notably, this acute timeframe has been overlooked by prior studies, which focused on long term (4+ weeks post-exposure) effects.^24–28^ We show that ZIKV infection and systemic stimulation with Poly(I:C) each induced significant changes in sperm miRNAs and tDRs within days of exposure (Figure 1B-C).^40^ The convergence of these responses implicates immune activation, independent of viral replication, is sufficient to remodel the sperm small RNA profile (Figure 1D).

To explore how the male reproductive tract responds to systemic inflammation in order to mediate small RNA alterations, we assessed gene expression via mRNA-seq in the testis and epididymis following Poly(I:C) stimulation. Interestingly, the epididymis, particularly the cauda epididymis, displayed a proinflammatory profile in response to Poly(I:C) stimulation (Figure 2, Supplemental Figure 2). We find that this tissue is potentially primed to respond to proinflammatory events due to expression of cytokine receptors, such as *Tnfrsf1a*, *Ifnar2*, and *Ifngr1,* under both basal and PIA induced conditions. Conversely, the testis was relatively unresponsive to Poly(I:C) stimulation, and expressed cytokine receptors at a lower level, highlighting the immune privileged nature of this tissue.^41^ We speculate that cytokine signaling is responsible for creating a proinflammatory environment in the epididymis (Figure 2), which ultimately changes the expression of small RNAs within the tissues, leading to differential accumulation of small RNAs within sperm (Figure 1), consistent with soma-to-germline transfer of RNAs via epididymosomes.^8–11^ Future studies are needed to define the role of specific cytokine signaling pathways, such as IL-6 and type I interferon signaling in mediating alterations to the sperm small RNA profile. By establishing a direct link between paternal immune sensing and epigenetic remodeling in sperm, our work provides a mechanistic framework for how the immune systems shapes epigenetic information in sperm.

Given this rapid remodeling of sperm small RNAs, we next considered the developmental consequences of PIA on early embryonic gene expression. We found that PIA leads to a robust reprogramming of the gene expression landscape in pre-implantation embryos (Figure 3). These findings demonstrate that PIA shapes offspring outcomes at the earliest stages of development. Remarkably, microinjections of sperm RNAs from PIA males delivered to naive embryos were sufficient to recapitulate the embryonic gene expression reprogramming observed following fertilization by PIA sperm (Figure 5). These findings demonstrate that altered small RNAs within the sperm of our PIA males causally modulate embryonic gene expression, providing a mechanistic link between paternal small RNAs and offspring phenotypes.

This study is the first to report sex-based differences in the context of intergenerational non-genetic inheritance observed during early embryonic development. We found that PIA induces sex specific transcriptional programs at multiple stages of embryonic development (Figure 4). Importantly, these changes are not linked to altered expression of genes on sex chromosomes (Figure 4, Supplemental Figure 3), suggesting that the embryo sex does not influence altered gene expression patterns. Rather, we speculate that these sex-based differences start at the preconception level in sperm, potentially through differential small RNA accumulation in sperm containing X vs Y chromosomes. In fact, it has previously been reported that X chromosome containing sperm express surface proteins that are not found on Y containing sperm.^42,43^ These distinct surface protein profiles may result in differential interactions with epididymosomes and thus altered small RNA cargo delivery in X and Y sperm. Though differential small RNA profiles in X and Y sperm have been reported in bulls,^44^ more work is needed to assess this phenotype in mice and explore the interactions between epididymosomes and sperm surface proteins.

While other PIA studies have focused primarily on offspring behavior,^24–26^ we specifically explored the impacts of PIA on offspring infection outcomes, an area that has remained largely unexplored.^28^ Previous work has shown that *C. albicans* infection in male parents alters the sperm methylome and confers resistance to bacterial infection in offspring.^28^ However, an additional study implementing multiple PIA and offspring infection models was unable to confirm these results.^29^ Importantly, these studies focus on offspring outcomes 4 weeks after paternal stimulation. In contrast, we specifically assessed how acute PIA altered offspring susceptibility to a lethal dose of influenza A. We found male offspring sired from acute PIA males showed dramatically better survival (∼71%) compared to female siblings and control sired offspring (∼15-18%) (Figure 6), with no link between survival and better control of viral replication or enhanced viral clearance from the lung (Supplemental Figure 5). These striking outcomes demonstrate the beneficial protection inherited by offspring of PIA males. Future studies should define the mechanism of protection and the breadth of protection to other microbial pathogens. Embryonic gene expression analysis further indicates that PIA begins to reprogram offspring immunity shortly after fertilization with immune related pathways and genes altered in both 4-cell and morula stage embryos (Figure 3). These findings suggest that PIA-induced offspring protection is conferred at the time of conception and retained throughout early life. How this protection is established and propagated throughout development warrants further investigation.

This study provides an extensive framework for how paternal immune activation shapes inheritance across generations, linking gene expression remodeling in the male reproductive tract and altered sperm small RNAs to remodeling of embryonic development and inherited pathogen protection in offspring. In non-mammalian models, parental pathogen exposure is known to enhance offspring immunity, a phenomenon termed transgenerational immune priming (TGIP).^17^ Paternal TGIP is thought to be mediated by epigenetic information, potentially small RNAs, drawing a direct parallel to non-genetic inheritance in mammals.^17–19^ Together with prior work, our findings here suggest that this form of heritable immunity is evolutionarily conserved across mammals, birds and insects. Epidemiological evidence further indicates that immune activation in humans, through vaccination, increases offspring survival in early life.^45^ Collectively, these observations establish non-genetic immune inheritance as a physiologically meaningful mechanism by which paternal immune experiences shape immune phenotypes in offspring.

## ACKNOWLEDGEMENTS

This work was in part supported by the Pew Charitable Trusts Biomedical Scholars Program (KAJ & CC) and Burroughs Wellcome Foundation Next Generation Pregnancy Grant (KAJ). Additional funding provided by the NSF GRFP (TME and ADC), the University of Pennsylvania Cell and Molecular Biology Training Grant T32 GM-07229 (MSM), HHMI Gilliam Fellowship (MCA), Eunice Kennedy Shriver National Institute of Child Health and Human Development F31HD114433 (GSL), Loan repayment program: Conception and Infertility Research L50HD119810 (GSL), and Hartwell Foundation Individual Biomedical Research Award (EAK).

## AUTHOR CONTRIBUTIONS

TME: study conceptualization, study design, methodology, experimental work, data interpretation, manuscript writing. ADC: methodology, experimental work, manuscript review and editing. EAK: experimental work, manuscript review and editing. MNL: experimental work, manuscript review and editing. CNM: experimental work, manuscript review and editing. GSL: methodology, experimental work, manuscript review and editing. MW: experimental work, manuscript review and editing. MCA: experimental work, manuscript review and editing. MSM: experimental work, manuscript review and editing. SGN: experimental work, manuscript review and editing. PH: experimental work. CCC: funding acquisition, study conceptualization, study design, methodology, manuscript writing, resources and supervision. KAJ: funding acquisition, study conceptualization, study design, methodology, manuscript writing, resources and supervision.

## RESOURCE AVAILABILITY

### Lead Contact

Requests for further information and resources should be directed to and will be fulfilled by the lead contacts, Colin Conine and Kellie A. Jurado.

### Materials Availability

This study did not generate new unique reagents.

### Data and Code Availability

- Sequencing data have been deposited at National Center for Biotechnology Information: accession number and are publicly available as of the date of publication.
- All original code has been deposited at https://doi.org/10.5281/zenodo.18851786 and is publicly available as of the date of publication.
- Any additional information required to reanalyze the data reported in this paper is available from the lead contacts upon request.

## METHOD DETAILS

### Mice

Human *STAT2* knock-in (hSTAT2KI) mice^32^ were obtained from the Diamond laboratory at University of Washington in St. Louis. *Infar1* knock out (IFNAR -/-) mice and wild-type C57BL/6J (B6) mice were procured from Jackson Laboratory. All mice were maintained at the University of Pennsylvania under specific-pathogen free conditions, following the University of Pennsylvania approved IACUC protocol (#806749).

### Virus

Zika virus DAKAR mouse-adapted (ZIKV) strain^32^ was obtained from the Diamond laboratory at the University of Washington in St. Louis. ZIKV was propagated in VERO cells and concentrated using ultrafiltration. VERO cells were grown in Dulbecco’s Modified Eagle’s Media (DMEM) supplemented with 10% heat-inactivated Fetal Bovine Serum (FBS) and 1% penicillin/streptomycin. Influenza A virus H1N1 strain A/Puerto Rico/8/1934 (influenza A) was obtained from the Hensley Laboratory at the University of Pennsylvania.

### Murine PIA Models

To model PIA using ZIKV, male IFNAR -/- and hSTAT2KI mice at 11-17 weeks old were infected with 1×10^5^ or 1×10^6^ plaque forming units (PFUs) of ZIKV DAKAR via footpad injection. Mice were monitored daily for changes in weight. Mice with a 20% reduction in weight were humanely euthanized. Spleen, testis, and epididymal tissues were collected at 5-, 7-, 10-, and 20-days post-infection into TRIzol or DMEM supplemented with 2% heat-inactivated FBS and homogenized. Sperm was collected from hSTAT2KI males at 5 days post-infection. To model PIA using Poly(I:C), 13-15 week old B6 mice were treated with 20mg/kg of High Molecular Weight (HMW) Poly(I:C) via intraperitoneal injection. Serum was collected at 6 hours post-treatment to confirm stimulation by assessing IL-6 induction. Testis and epididymal tissues and sperm samples were collected at 48 hours post treatment into TRIzol and homogenized.

### Plaque Assay

Testicular and epididymal tissue homogenates collected into DMEM supplemented with 2% heat-inactivated FBS were used to assess Zika viral burden by plaque assay. Homogenates were serially diluted in DMEM. Vero cells were seeded at 600,000 cells per well in 6-well plates for plaque assay. 24 hours post-seeding, Vero cells were inoculated with 250μl of diluted sample homogenate and incubated for 1 hour at 37°C. Plates were rotated in a figure 8 motion every 15 minutes during this incubation. Following the incubation, samples were aspirated from the wells and covered with an overlay comprised of Minimum Essential Medium (MEM) supplemented with 5% heat-inactivated FBS, 1% GlutaMAX, 1% non-essential amino acids, and 0.65% agarose. At 6 days post-infection, cells were fixed with 10% Neutral Buffered Formalin (NBF) and cells were stained using 0.1% Crystal Violet. Individual plaques were counted and viral titer was calculated.

### RNA isolation, cDNA synthesis, and quantitative RT-qPCR

Testicular, spleen, and epididymal homogenates harvested into TRIzol reagent were used to assess ZIKV viral burden by RT-qPCR. 100μl of sample was diluted in an additional 300μl of TRIzol reagent and mixed with 400μl of 100% ethanol. RNA was isolated following the manufacturer’s instructions for the ZYMO Direct-zol RNA miniprep kit. cDNA was synthesized from isolated RNA following manufacturer’s instruction for the iScript cDNA synthesis kit. Quantitative RT-qPCR was performed using SYBR Green Mastermix reagent using the following ZIKV specific primers: Forward 5′-CCACCAATGTTCTCTTGCAGACATATTG-3′; Reverse 5′-TTCGGACAGCCGTTGTCCAACACAAG-3′.^32^ ZIKV equivalents were calculated using a 10-fold serial dilution standard curve isolated from viral stock. ZIKV equivalents were standardized to tissue weight in mg.

### Sperm isolation

For sperm isolation, Whitten’s medium was made by combining 100mM NaCl, 4.7mM KCl, 1.2mM KH_2_PO_4_, 2.3mM MgSO_4_, 5.5mM Glucose, 1mM Pyruvic Acid, 4.8mM Lactic Acid, 20mM HEPES in Nuclease Free Water (NFW). To isolate mature sperm, cauda epididymides were dissected and placed into a dish with 1.5mL of Whitten’s medium. Sperm were removed from the cauda by gently squeezing caudal lumen contents into the media. Sperm were left at 37°C for 10 minutes and then transferred to a microcentrifuge tube. Sperm were kept at 37°C for 5 minutes to allow separation of sperm and debris. Supernatant containing the sperm was transferred to a new tube and sperm were pelleted by spinning at 10,000 x g for 5 minutes. Sperm pellets were washed twice with PBS and treated with somatic cell lysis buffer (0.01% SDS and 0.005% Triton-X in NFW) for 10 minutes to remove remaining somatic cells. Sperm were washed with PBS an additional time before RNA isolation.

### RNA isolation

To isolate RNA from sperm, sperm were resuspended in sperm lysis buffer (6.4 M Guanidine HCl, 5% Tween-20, 5% Triton, 120mM EDTI, and 120mM Tris pH 8.0), 0.66mg/mL Proteinase K, and 33mM DTT. Sperm were incubated at 60°C for 15 minutes. One volume of NFW and 2 volumes of TRIzol reagent was added to the sample after incubation and mixed well. Samples were added to prespun phase lock tubes with 0.2 volume of 1-bromo-2-chloropropane (BCP reagent) and mixed by inversion 15 times. Tubes were spun at 18,000 x g for 4 minutes at 4°C. The top layer was transferred to a low bind microcentrifuge tube with 1μl of GlycoBlue Coprecipitant and 1.1 volume of isopropanol and mixed well. Samples were allowed to precipiate for at least 1 hour at -20°C. RNA was pelleted by spinning at 18,000 x g for 15 minutes at 4°C. RNA was washed and repelleted in cold 70% ethanol and reconstituted in NFW. To isolate RNA from tissues for small RNA or mRNA sequencing, homogenized tissue samples in TRIzol were transferred to pre-spun phase lock tubes. 0.2 x volume of BCP reagent was added to each tube and mixed by inversion (15 times). Tubes were spun and RNA precipitation was carried out as described above. RNA from tissues was DNase treated with 10μl of RDD buffer and 2.5μl of DNase I and incubated at room temperature for 15 minutes. 100μl of NFW and 400μl of TRIzol reagent were added to each sample. Samples were added to pre-spun phase lock tubes with 120μl of BCP reagent and mixed by inversion. Samples were spun and precipitated as previously described.

### Small RNA isolation and sequencing

For small RNA isolation from sperm, the entirety of the samples was used for isolation. For small RNA isolation from tissue, 200ng of RNA was used for isolation. The small RNA fraction was isolated from each sample by running samples on a 15% polyacrylamide-7M urea denaturing TBE gel. Size selection was used to isolate the 18-40 nucleotide region (small RNA fraction). Small RNA libraries were prepared using a modified Illumina Tru-Seq small RNA library prep protocol. Briefly, T4 truncated ligase was used to ligate an adapter to the 3’ end of small RNAs and 5’ end adapters were added with T4 ligase. cDNA was synthesized using Superscript II reverse transcriptase kit. cDNA was amplified via qPCR to generated individual libraries. A subset of samples, representing each tissue and input amount were used to determine the minimum number of amplification cycles necessary. Libraries were gel-purified, quantified, and pooled for sequencing on a NextSeq 1000. Read quality was assessed using FastQC. Adapter sequences were trimmed from reads using trimmomatic and reads were sequentially mapped to rRNA mapping reads, miRbase, murine tRNAs, pachytene piRNA clusters, repeatmasker, and Refseq using Bowtie 2. These bioinformatic processes were conducted through a pipeline hosted on DolphinNext.^46^ Mapped reads were normalized to total genome reads minus rRNA reads.

### mRNA-sequencing

For mRNA-sequencing on tissue samples, 75-500ng of sample was used to construct mRNA-sequencing libraries using the Illumina stranded mRNA kit. Briefly, messenger RNA was captured using oligo(dT) magnetic beads, fragmented, and primed for cDNA synthesis. Synthesized cDNA was cleaned using AMPure XP beads and libraries were amplified via PCR. A subset of samples, representing each tissue and input amount were used to determine the minimum number of amplification cycles necessary. Libraries were quantified using a Qubit and were combined at equal concentrations. The final combined library was purified using non-denaturing PAGE. Pair-end sequencing was performed on a NextSeq 1000. On a bioinformatics pipeline hosted on DolphinNext,^46^ read quality was assessed using FastQC and reads were mapped to the *Mus musculus* genome (mm10) using RSEM. Reads were normalized to transcripts per million. The DESeq2^47^ package in R was used to determine differentially expressed genes (DEGs). DEGs were defined by fold-change > 1.5 and adjusted p-value ≤ 0.05. GO analysis was performed by feeding the top 50 up- or down-regulated genes for each tissue into GProfiler.^48^ g:GOSt was used to query genes against the *Mus musculus* dataset with default parameters.

### In vitro fertilization and embryo culture

5-week-old female B6 mice were superovulated for egg collection. Briefly, 5IU Pregnant Mare Serum Gonadotropin (PMSG) was administered via IP injection. 48 hours later, 5IU Human Chorionic Gonadotropin (hCG) was administered via IP injection. 12-15 hours post-hCG treatment, ovaries were removed. Cumulus-oocyte complexes were removed from the oviductal ampullae and moved to warm human tubal fluid (HTF) media. Complexes were moved to HTF supplemented with 1.0mM reduced glutathione (GSH) for in vitro fertilization (IVF). Oocytes were divided into two groups to be fertilized by sperm from control or PIA males. Male mice were treated with Poly(I:C) or PBS as described above. Sperm were collected 48 hours post treatment from the cauda epididymis as described above into Biggers Whitten and Whittingham (BWW) media ((91.5 mM NaCl, 4.6 mM KCl, 1.7 mM CaCl_2_·2H_2_O, 1.2 mM KH_2_PO_4_, 1.2 Mm MgSO_4_·7H_2_O, 25 mM NaHCO_3_, 5.6 mM D-glucose, 0.27 mM sodium pyruvate, 44 mM sodium lactate, 5 U/mL penicillin, 5 μg/mL streptomycin, 20 mM HEPES buffer, and 3.0 mg/mL bovine serum albumin (BSA)) supplemented with 1 mg/mL methyl-β-cyclodextrin. Sperm were capacitated for 45 minutes at 37°C under an atmosphere of 5% O_2_, 6% CO_2_. 2×10^5^ capacitated sperm were added to IVF drops (HTF supplemented with GSH containing oocytes). IVF drops were incubated for 3 hours at 37°C under an atmosphere of 5% O_2_, 6% CO_2_. Zygotes were washed through KSOM droplets following incubation and allowed to develop in fresh KSOM droplets. Embryos were collected at the 4-cell (46 hours post-fertilization) and morula (72 hours post-fertilization) developmental stages.

### Single embryo RNA injections

Sperm was collected from six Poly(I:C) treated males and RNA was isolated as described above. RNA was reconstituted in one tube. RNA was DNase treated as described above. 400ng of RNA was retained for total RNA injections. Remaining RNA was used to isolated small RNAs by size selection in a 5% polyacrylamide-7M urea denaturing TBE gel as described above. Total RNA was diluted to 50ng/μl and frozen for injections. Small RNAs were diluted to 4ng/μl and frozen for injections. Small RNAs were diluted to 2ng/ul for injections on the day of the procedure.

Zygotes for injections were generated by fertilizing eggs from naive superovulated female mice with sperm from control males as described above. After fertilization, zygotes were washed in KSOM. Zygotes were transferred to injection plates in FHM with 0.001% PVA droplets. RNA injections were performed using a Femtojet microinjector with Femtotip II microinjection capillary tips set to 100 hPa pressure for 0.2 seconds with 7 hPa compensation pressure. After injection, zygotes were returned to KSOM droplets and allowed to develop as described above.

### Single embryo mRNA isolation and sequencing

Individual embryos at the 4-cell and morula developmental stages were collected into 1 x TCL buffer supplemented with 1% 2-mercaptoethanol for embryo cell lysis. Libraries for sequencing were generated with the SMART-Seq protocol. Briefly, RNA was isolated using RNAClean-XP beads and reverse transcribed using the Superscript II kit. 10 cycles were used to amplify cDNA using indexes from the Nextera XT kit. Libraries were sequenced using a NextSeq 1000. Using our previously described platform on DolphinNext,^46^ reads were mapped to the *Mus musculus* genome (mm10) and normalized to transcripts per million. DEGs were determined using DESeq2^49^ as previously described with a threshold of fold-change ≥ 1.5 and p-value ≤ 0.01. Ingenuity pathway analysis was performed on DEG lists via Qiagen’s online platform.^50^ DEG chromosome location was determined using MGIBatch Query^51–53^ using the C57BL/6J genome location option under gene attributes. Venn diagrams to visualize overlapping DEGs were made using InteractiVenn.^54^

### Offspring generation

To generate offspring, 13-week-old male mice were treated with Poly(I:C) or PBS as described above. Serum was collected at 6 hours post-treatment to confirm immune stimulation by IL-6 ELISA. 40-42 hours post-treatment, females were placed with treated males to mate overnight. The following morning, females were checked for the presence of a copulation plug and all females were removed from the cages with treated males. Successfully plugged females were separated from unmated females. At embryonic day (E) 14.5 or 15.5 (E14.5 or E15.5), pregnant females were singly housed for pup delivery. At E19.5, successful pup delivery was confirmed by visual inspection. Offspring were weaned at 21 days old and allowed two days to adjust to new cages and dam removal prior to infection.

### Influenza A infection

Influenza A PR8 virus was diluted to 250 TCID50 in PBS. Mice were sedated using isoflurane and were given 25μl of inoculum intranasally (12.5μl per nostril). Mice were monitored out to 14 days post-infection, with weight, rectal (core) temperature, and clinical score recorded daily. Clinical signs are defined as piloerection, hunched posture, dyspnea, and absence of escape response. Mice that reached a 30% reduction in weight, temperature below 30°C, or displayed all four clinical signs were humanely euthanized.

### TCID50 assay

To assess influenza viral burden, right lung lobes were harvested from infected mice at 4- or 6/7-days post-infection. Lungs were harvested into serum free MEM and homogenized. MDCK cells were seeded at 50,000 cells/ well in 96-well plates. MDCK cells were maintained in MEM supplemented with 10% heat-inactivated FBS. 24 hours post-seeding, plates were washed twice with 200μl warm MEM and 180μl of TCID50 medium (50mL MEM, 50μl Gentamycin sulfate, 50μl TPCK-trypsin, 250μl HEPES) was added to each well. Homogenized samples were spun at 8,000 rpm for 3 minutes and 20μl of supernantant was added to the first column of the 96 well plate with infection medium in quadruplicate. 10-fold dilutions were performed across the plate, with the last well remaining as a control. Plates were incubated at 37°C. After 3 days, plated were removed as individual wells were visually evaluated for cytopathic effect (CPE). Reed and Muench method was used to calculated TCID50/mL titer for each sample.

### IL-6 ELISA

Blood samples were collected 6 hours post-Poly(I:C) or PBS treatment via submandibular vein puncture. Samples were collected into Micro sample tubes with Serum Gel. Blood was allowed to clot for 30 minutes before tubes were spun at 10,000 x g for 5 minutes. Serum was transferred to clean microcentrifuge tubes and stored at -80°C prior to IL-6 ELISA. Stripwell Microplates were coated with anti-IL6 capture antibody (diluted 1:1000) and incubated at 4°C overnight. The following morning, plates were washed with ELISA wash buffer (0.05% Tween 20 in PBS) three times. Plates were incubated with blocking buffer (1% BSA in PBS) for 1 hour at room temperature. Plates were washed again with wash buffer three times then samples, diluted 1:50 in blocking buffer were added to the plate and incubated at room temperature for 2 hours. Incubation was followed with a wash using wash buffer three times before the addition of biotin anti-IL-6 detection antibody (diluted 1:1000) and incubated at room temperature for 1 hour. Another round of washes was performed before the addition of HRP-streptavadin (1:1000 dilution), which was incubated at room temperature for 30 minutes. A series of five washes using wash buffer was performed with the buffer sitting in the wells for 30 seconds prior to removal during each wash. An equal parts mixture of TMB substrate A and substrate B was added to the plate for 15 minutes and then stop solution was added. Plates were read using a Spectromax plate reader at 540nm with a wavelength correction of 450nm.

### Figure Creation

Figures 1A, 2A, 3A, 5A, and 6A were created using BioRender. Final edits to figures 2D, 2E, 3C, 3E, 4A, 4B, 4D, and 4E were completed in Adobe Illustrator after generation in R.

