## Supplemental Figures for "Acute Paternal Immune Activation Shapes Embryonic Development and Protects Offspring from Viral Infection"

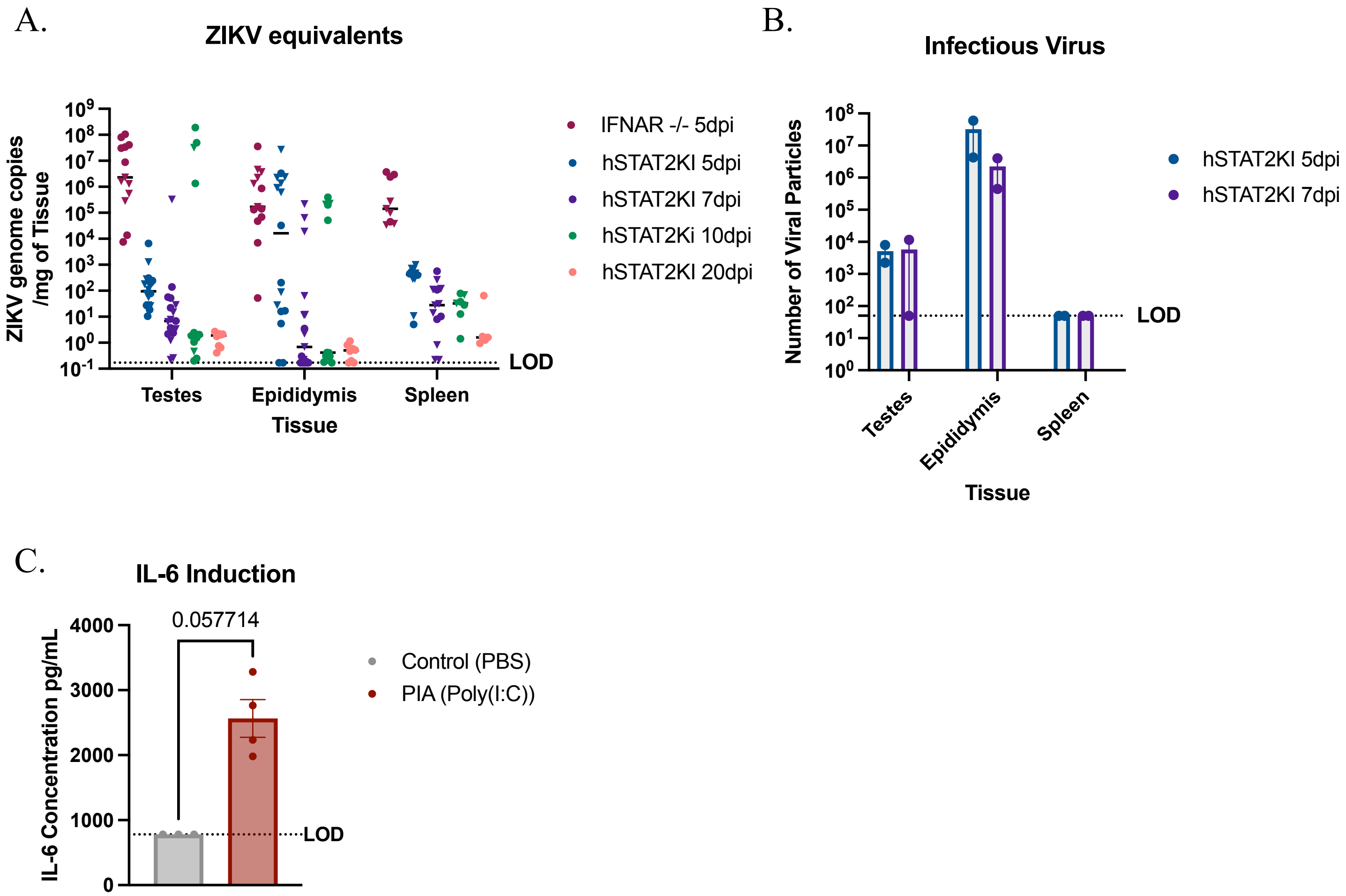


**Figure S1.** Male IFNAR -/- and hSTAT2KI mice were infected with 10^6 PFUs ZIKV DAKAR MA strain. Tissues were collected at 5-, 7-, 10-, and 20-days post infection. Viral burden was measured by qPCR (**A.**) and plaque assay (**B.**). **C.** 13-week-old male mice were treated with Poly(I:C) and 6 hours later, serum was collected to assess IL-6 induction by ELISA. Mann-Whitney Test.

A.

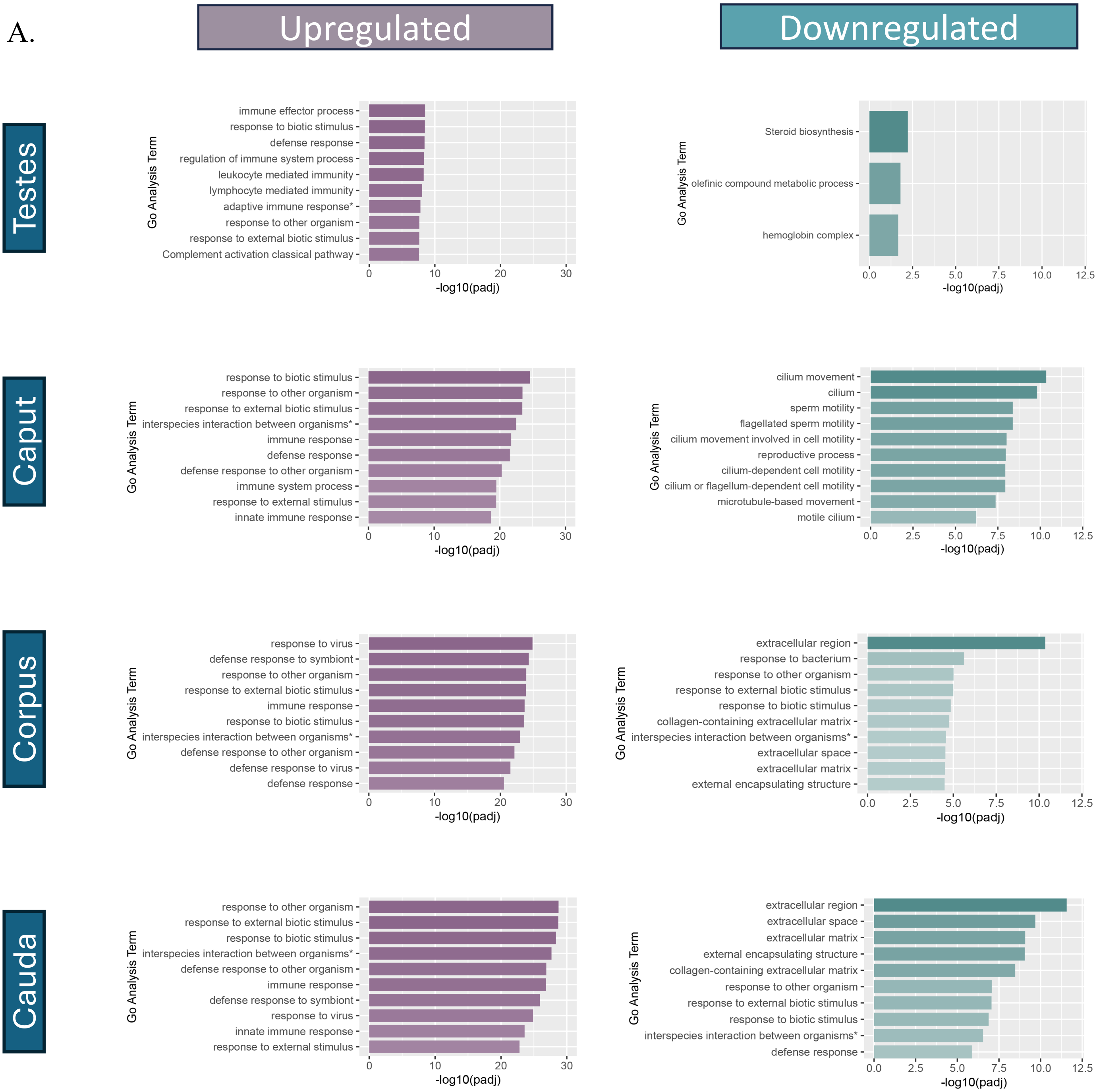


**Figure S2. Poly (I:C) treatment alters expression of immune related pathways primarily within the epididymis. B.** The top 50 up- and downregulated genes in each tissue were used for GO analysis using GProfiler. The top 10 up- and downregulated GO analysis terms are shown by adjusted p value. Adaptive immune response* = adaptive immune response based on somatic recombination of immune receptors built from immunoglobulin superfamily domains. Interspecies interaction between organisms* = biological process involved in interspecies interaction between organisms.

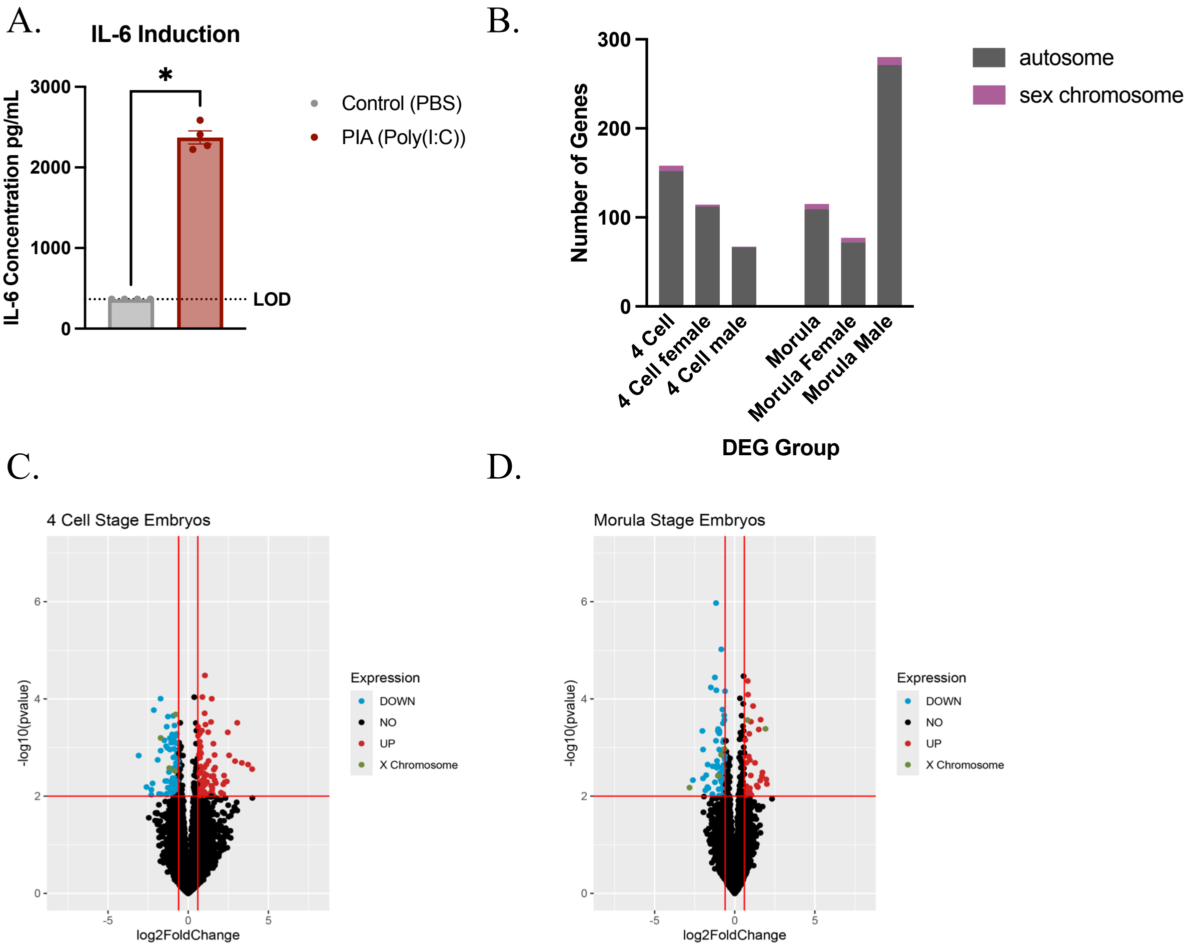


**Figure S3. Changes in early embryonic gene expression when fertilized with sperm from PIA fathers are not sex chromosome dependent. A.** 13-15-week-old male mice were treated with Poly (I:C). At 6 hours post-treatment, serum was collected and circulating IL-6 levels were measured by ELISA. Mann-Whitney Test, p-value = 0.0286. **B.** Number of differentially expressed genes (DEGs) in each group encoded on autosomes or the X chromosome. No DEGs were located on the Y chromosome. Chromosome location was determined using MGI Batch Report. **C.** Volcano plot showing differential gene expression in 4-cell stage embryos. **D.** Volcano plot showing differential gene expression in morula stage embryos. **C and D.** Red lines indicate significance thresholds ($-0.58 > \log_2FC$ or $\log_2FC > 0.58$ and p value $< 0.01$). Blue dots = downregulated, red dots = upregulated, green dots = genes encoded on the X chromosome.

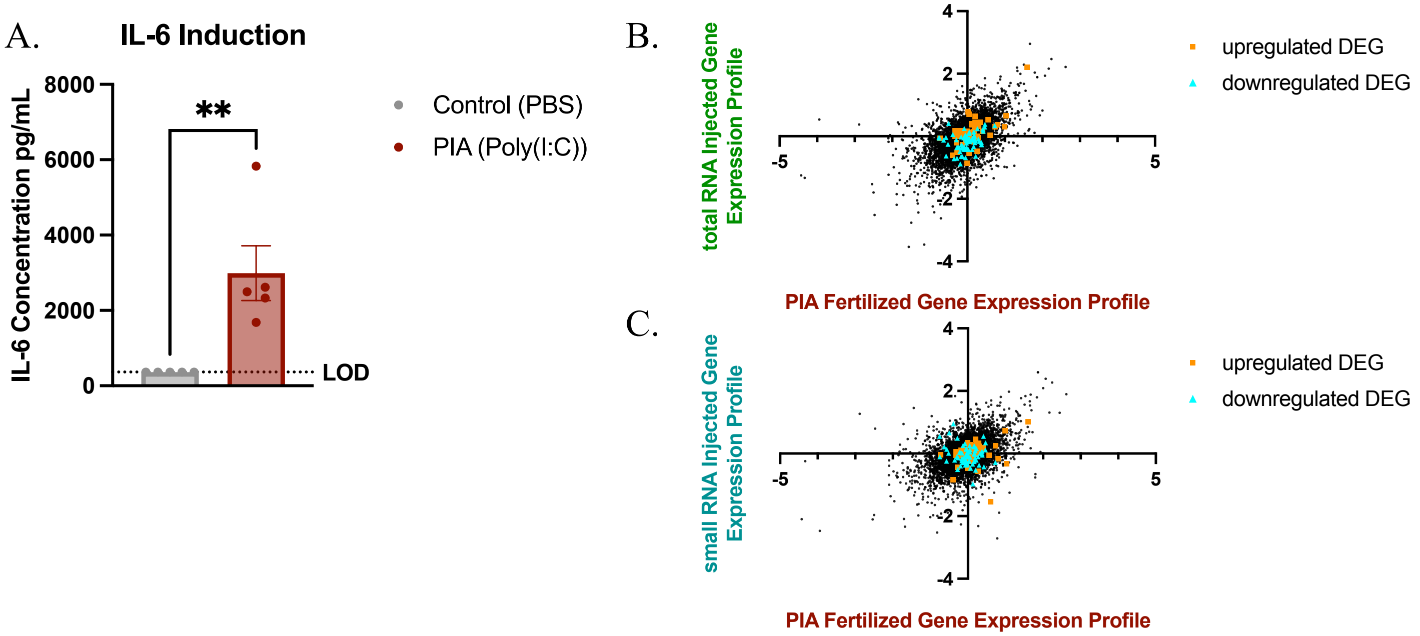


**Figure S4. A.** 13-15-week-old male mice were treated with Poly (I:C). At 6 hours post treatment, serum was collected and circulating IL-6 levels were measured by ELISA. Mann-Whitney Test, p-value = 0.0079. **B.** Comparison of profile established by total RNA injection to PIA fertilized profile, both profiles normalized to NFW embryos. Pearson Correlation. r = .5398 P-value <0.0001. **C.** Comparison of profile established by small RNA injection to PIA fertilized profile, both profiles normalized to NFW embryos. Pearson Correlation. r = .4391 P-value <0.0001. **B and C.** Orange squares indicate genes that were upregulated in original embryo sequencing experiments (Figure 3). Blue triangles indicate genes that were downregulated in original embryo sequencing experiments (Figure 3). All significant genes from Figure 3 are displayed.

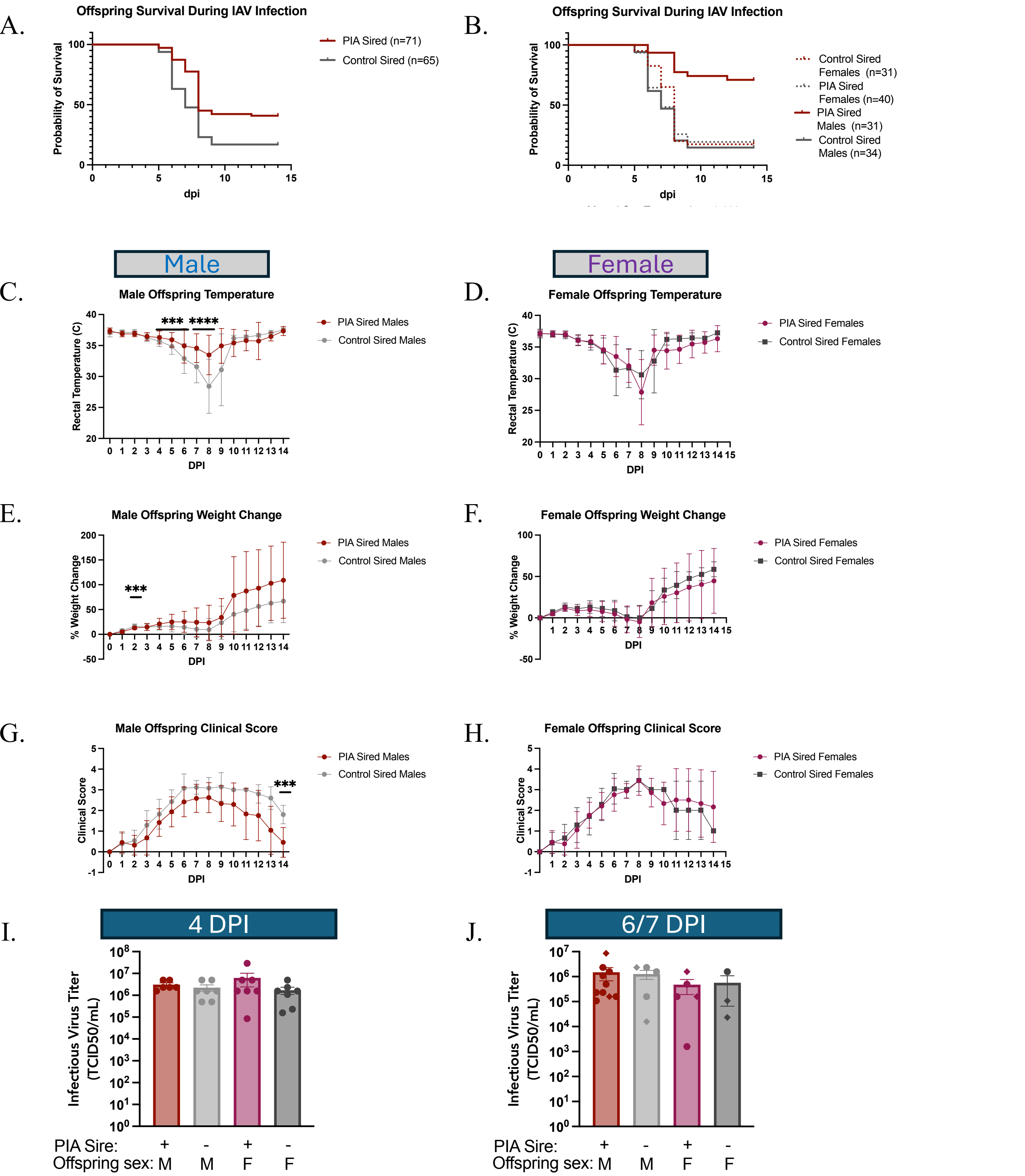


**Figure S5. PIA protects male offspring from negative outcomes during IAV infection. A.** Survival of offspring based upon sire condition. Mantel Cox Test: p-value = 0.0001. **B.** Survival of male and female offspring by sire condition. Mantel Cox Test: p-value <0.0001. Mice were monitored daily during infection for changes in temperature (**C and D**), changes in weight (**E and F**), and clinical signs of infection (**G and H**). Multiple Mann-Whitney Tests. *** = p-value <0.001, **** = p-value < 0.0001. Lungs were harvested at 4 (**I**) or 6/7 (**J**) days post infection and viral burden was measured by TCID50.
